# Joint inference of transcription factor activity and context-specific regulatory networks

**DOI:** 10.1101/2022.12.12.520141

**Authors:** Chen Chen, Megha Padi

## Abstract

Transcriptional regulation is a critical process that determines cell fate and disease. One of the challenges in understanding transcriptional regulation is that there is no easy way to infer the main regulators from gene expression data. Many existing methods focus on estimating the activity of individual transcription factors (TFs) using static TF-gene interaction databases, but regulomes are often altered in different cell types and disease conditions. To address this problem, we developed a new algorithm – Transcriptional Inference using Gene Expression and Regulatory data (TIGER) – that leverages Bayesian matrix factorization to simultaneously infer TF regulomes and transcription factor (TF) activities from RNA-seq data. We show that, when applied to yeast, A375, and MCF7 TF knock-out datasets, TIGER can provide more accurate predictions than comparable methods. The application to single-cell RNA-seq data reveals TIGER’s potential for uncovering cell differentiation mechanisms. Our results reinforce the importance of incorporating context-specific regulation when studying the mechanisms driving disease in different cell types.

## Introduction

A critical aspect of systems biology is to understand transcriptional regulation, i.e., to identify regulators driving a particular function and to determine the activities of those transcription factors (TFs) [1]. However, the challenge is that TF activity is difficult to measure, and the mRNA expression of a TF is not a reliable surrogate for its activity. Further, the targets of these TFs can change depending on the cell type, disease context, epigenetic state, and many other factors. Because of these problems, computational techniques have emerged to infer regulatory networks and the associated activity of the TFs.

Existing network inference methods can be divided into two categories by how much information they integrate. The first category is based solely on the gene expression data. One representative approach is the gene co-expression-based methods such as WGCNA [2], ARACNe [3], and Gaussian Graph Model [4]. These methods employ statistical analyses of dependencies between mRNA expression levels. In general, these methods can be used as an indicator for co-regulation; causality between genes can only be assumed if regulator genes are known in advance. The second category of methods incorporates TF binding information with gene expression datasets to infer a high-quality genome-scale regulatory network. These methods work because physical evidence of regulatory interactions (e.g., TF binding) and functional evidence (e.g., coordinated expression) provide complementary information for gene regulation [5]. For example, PANDA [6] applies network fusion to gene expression and sequence motif data to reconstruct genome-wide regulatory networks. TF binding data can be accessed through motif or ChIP-seq databases, such as JASPAR [7], TRANSFAC [8], RegulonDB [9], etc. This TF binding information can also be used to filter the reverse-engineered co-expression network. Applying this principle to single cell data, SCENIC [10] uses motif or ChIP data to filter out false positive edges in the GENIE3 [11] network. Inferelator 3.0 [12] incorporates scATAC-seq data for regulatory network inference. Comprehensive reviews of other network inference algorithms can be found in [13, 14].

Algorithms have also been developed to estimate TF activities (TFA) [15]. For instance, VIPER infers TFA from gene expression data based on each TF’s “regulon” (i.e., its target genes) and the sign of the regulatory interactions (i.e., activation or inhibition) [15]. These regulons can be defined in any way the user desires. However, studies have shown that VIPER performs better, on both bulk RNA-seq and single-cell RNA-seq data, when using literature-curated TF regulons rather than context-specific regulons inferred from data [16]. VIPER also considers every TF completely independent of each other and ignores potential multi-TF regulation of genes. Other methods have modeled the effect of multiple TFs and context-specific regulons. For example, Inferelator-AMuSR [17] first uses linear regression to infer TFA from gene expression data and TF binding priors and then updates network edge weights using multitask regularized regression. Similarly, [18] uses bilinear regression to simultaneously estimate TFA and the regulatory network. However, these two methods have only been tested on data from model organisms. Recently, a single-cell TFA inference method – BITFAM [19] was developed to estimate regulatory network and TFA using prior network and scRNA-seq data. However, it does not distinguish between TF activation and inhibition and it was not validated using gold standard data (e.g. TF knockout or overexpression data).

Here, we introduce a new method called TIGER (Transcriptional Inference using Gene Expression and Regulatory data) that jointly infers a context-specific regulatory network and corresponding TF activities. TIGER is inspired by many matrix factorization methods for TF or pathway activity analysis, such as Network Component Analysis (NCA) [20], PLIER [21], Sabatti et al. [22], and Dai et al. [23]. These methods range from frequentist to full Bayesian matrix factorizations. They implement network sparsity and provide edge weight estimates but do not distinguish between activation and inhibition of transcription. Therefore, for TIGER, we developed a new Bayesian framework that can incorporate both sparsity and sign constraints by imposing tailored prior distributions on the variables. In addition, we implement a Variational Bayesian method [24, 25] to accelerate the estimation process. When applied to yeast, human cancer cell lines, and scRNA-seq data, TIGER can more sensitively infer cell-type-specific TF activities than comparable methods. TIGER’s Bayesian framework enables data-driven learning of context-specific signed regulons for accurate estimation of regulatory activity in mammalian and single-cell data.

## Results

### Overview of TIGER

TIGER estimates a gene regulatory network (GRN) and TF activity (TFA) levels through a matrix factorization framework (Figure 1A). Specifically, we aim to decompose an observed log-transformed normalized gene expression matrix *X* into a product of two matrices, *W* and *Z*, where *W* is interpreted as a regulatory network, and *Z* is interpreted as a TFA matrix. To guide the construction of biological meaningful GRN and TFA, we add both network structure and edge sign constraints to *W*, and constrain TFA to be strictly non-negative. The network structure curated from literature (e.g., ChIP-seq, TFBS) is used as a prior distribution to guide the GRN estimation. Non-negative constraints on TFA guarantee that the edge signs can be interpreted as activation and suppression events. We use a Bayesian framework to link prior knowledge and constraints to gene expression data. As the literature information for the prior network is usually very general, we use a sparse prior to filter some context-irrelevant edges. Thus, the posterior network *W* will be a context-specific network, and the estimated TFA will reflect the regulatory activities for each sample. See Methods and Supplementary Information for details of the TIGER algorithm.

**Figure 1.**
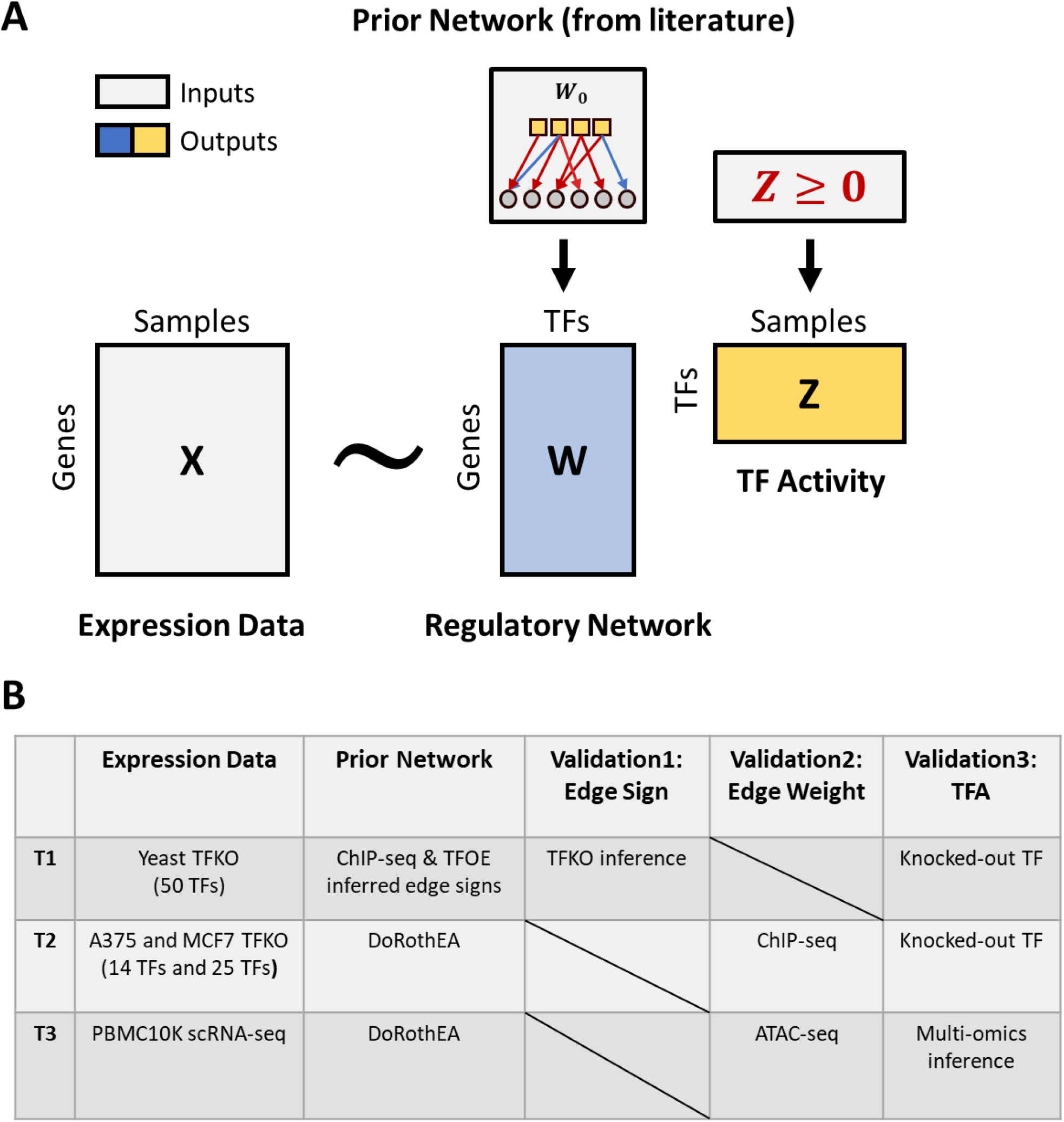
Overview of TIGER. (A). TIGER uses a matrix decomposition method to estimate regulatory network *W* and TF activity *Z*. The inputs of TIGER are normalized gene expression matrix *X* and prior TF binding network *W*_0_ curated from the literature. Prior binding information is incorporated as different prior distributions in matrix *W*. Matrix *Z* is constrained to be strictly non-negative for identifiability and biological interpretation. Variational Inference is used to estimate parameters. (B). Description of the validation strategy. Test 1 (T1) uses the Yeast TFKO dataset and ChIP-seq data to assess the impact of sign-flipping. Test 2 (T2) uses A375 and MCF7 cancer cell line TFKO datasets and the DoRothEA prior network to assess the importance of edge weights. Test 3 (T3) explores TIGER’s potential in scRNA-seq data analysis.

To test the TIGER model, we use three TF knock-out (TFKO) datasets and one scRNA-seq data (Figure 1B). Since TFA is not easy to measure in the lab, we use TFKO datasets where the knocked-out TF should have the lowest activity in that sample. To validate the structure of the inferred GRN, the most often used gold standard is ChIP-seq data. In the first test (T1), we use gene expression data from 50 different yeast TF knockout strains and test whether TIGER can identify the knocked-out TF in each sample. In the second test (T2), we increase the complexity of the problem. First, we change the setting to cancer cell lines, namely A375 melanoma cells (14TFs knocked out) and MCF7 breast cancer cells (25 TFs knocked out), representing a much larger genome and more complex regulatory mechanisms. Second, we use a general prior network called “DoRothEA” [26]. The goal of T2 is not only to test TFA estimation but also the accuracy of the inferred GRN. We found that 11 TFs in the A375 and MCF7 TFKO cell lines have ChIP-seq data available in Cistrome DB [27]; thus, we can determine whether TIGER correctly assigns higher edge weights to the true edges. In the third test (T3), we apply TIGER to a highly used 10x multiome PBMC single-cell dataset publicly available on the 10x website. In this dataset, paired scRNA-seq and ATAC-seq profiles are measured in 10,412 PBMCs, and can be integrated in a multi-modal analysis to create a gold standard for the TFA. We used this to evaluate the inference of TFA from only the scRNA-seq data using TIGER.

As a comparison for TIGER, we applied VIPER [15], a well-established method for estimating TF activities using the aREA method, a statistical test based on the average ranks of the target genes. VIPER is an appropriate comparison method for TIGER because it is commonly used to estimate TFA and it takes in the same two inputs as TIGER – gene expression profiles and a signed prior regulatory network. An advantage of VIPER is that it can estimate TF activities for single samples, whereas TIGER needs multiple samples as input. A potential advantage of TIGER is that it can update and potentially improve the prior network, whereas VIPER relies on the accuracy of the prior. We applied both methods to all three test datasets and ranked the TFA estimates in each sample. In the TFKO datasets, the TF with the lowest estimated TFA would be ranked 1 because we are searching for the knocked-out TF. However, in the scRNA-seq data, our goal is to identify the driver TFs so we assign the TF with highest activity a rank of one.

### TIGER improves TF activity estimation by improving edge sign accuracy

We first tested TIGER’s performance on a Yeast TFKO dataset (T1). The prior network was curated from ChIP data [18]. The signs for the prior were assigned by calculating whether the target gene was increased or decreased in response to overexpression of that same TF in an independent dataset [18]. Both TIGER and VIPER perform very well on this dataset, with both methods correctly identifying 20 out of 50 TFs as rank No.1 in the corresponding knock-out sample (Figure 2A). TIGER is slightly better than VIPER when considering the average rank of all 50 TFs. To further evaluate the consistency between TIGER and VIPER, we first computed the Spearman correlation of their TFA rankings and found a high consistency of R=0.65 (p<0.05) between TIGER and VIPER (Figure 2B). If one defines a “success” as the knocked-out TF being among the top 10 results, TIGER and VIPER are both successful for 30 TFs and both fail for 9 TFs. Among the remaining 11 TFs, VIPER succeeds for 1 of them (and TIGER fails), but TIGER succeeds at 10 TFs (and VIPER fails).

**Figure 2.**
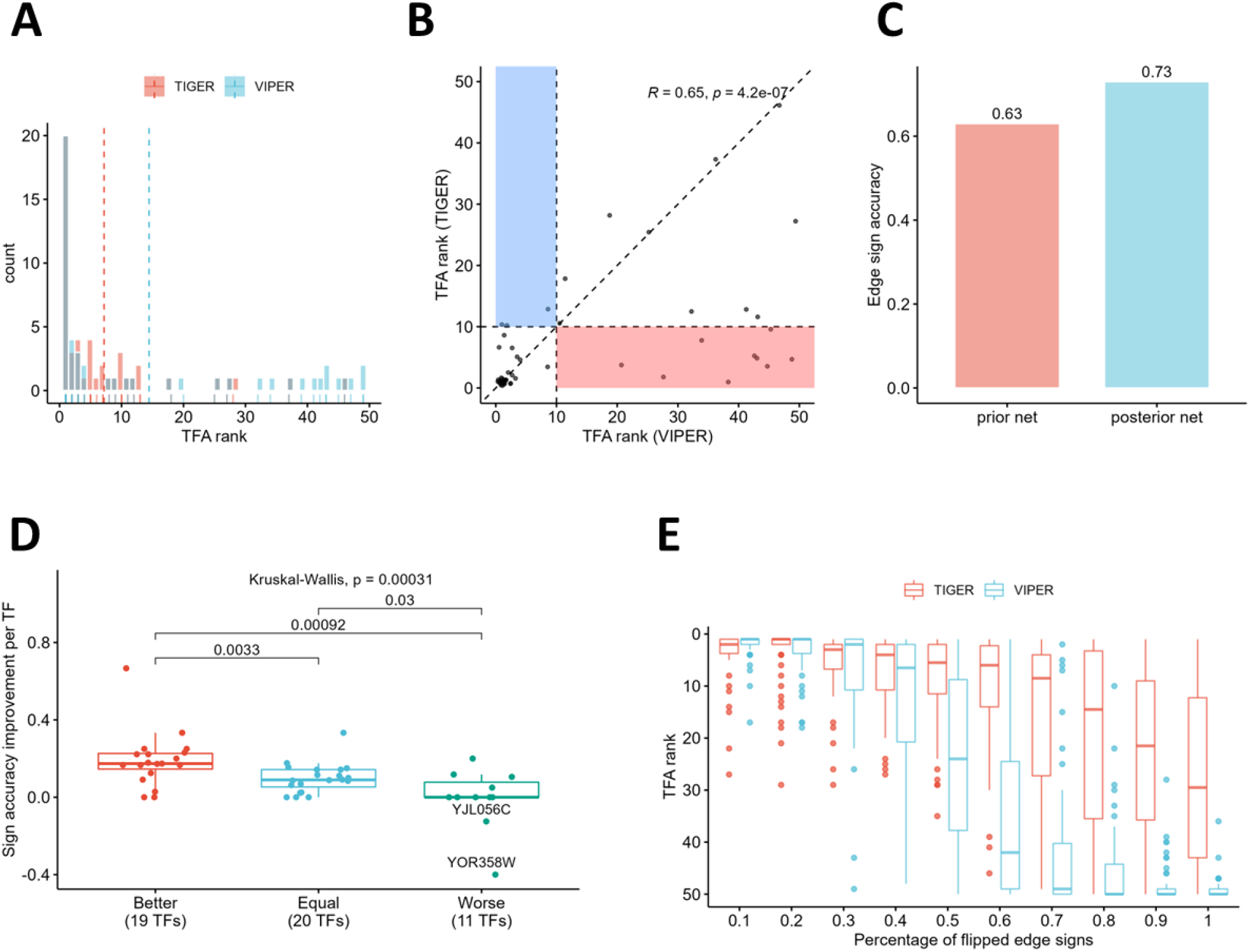
Comparison of TIGER and VIPER on yeast TF knock-out dataset. (A). Histogram of TIGER and VIPER’s ranking of knocked-out TFs. Dashed lines are the average ranking by each method. A lower rank is better. (B). Consistency between TIGER and VIPER on TFA estimation. Spearman correlation R = 0.65 (p<0.05). The red rectangular region covers the 9 TFs that TIGER successfully identifies, but VIPER fails. The blue rectangular region covers 1 TF that VIPER successfully identifies, but TIGER fails. (C). Bar plot of edge sign accuracy in the prior network and posterior network estimated by TIGER. (D). Boxplot shows the associations between edge sign accuracy improvement (Y-axis) and TIGER’s performance (X-axis). “Better,” “Equal,” and “Worse” stands for TFs for which TIGER has a better, equal, or worse prediction than VIPER. Kruskal-Wallis test p<0.05. Two TFs that have negative improvement are labeled. (E). Simulation results of random edge sign flipping in prior network.

We hypothesized that the cases in which TIGER has better performance than VIPER are driven by TIGER’s ability to improve the prior network. Because the prior network structure already uses high-quality ChIP data, we focused on evaluating changes in the edge signs. To define a “gold standard” for the edge signs, we used the change in target gene expression directly measured in the TF knockout data, rather than in the overexpression data that was used as input to the method. Comparing the edge sign accuracy for the prior and posterior network, we confirmed that TIGER improves this accuracy score from 0.63 to 0.73 (Figure 2C). Then we asked if TIGER outperforms VIPER by modifying edge signs. To answer this question, we separated all TFs into 3 groups – “Better,” “Equal,” and “Worse,” which correspond to TIGER having a better, equivalent, or worse ranking than VIPER. We computed the edge sign accuracy improvement for each TF, defined as the posterior accuracy minus prior accuracy. We found that TIGER’s performance is highly correlated with the edge sign accuracy improvement (Figure 2D, KW test, p<0.05).

Finally, we asked why TIGER ranks the true knocked-out TFs lower than VIPER in the “Worse” category. Examining these 11 TFs more closely, we noted that the edge signs for these TFs only have a tiny improvement, and in some cases edge sign accuracy is decreased. However, a large decrease in edge sign accuracy did not necessarily correspond to a large shift in TF ranking; for example, the TF YOR358W has the largest decrease of sign accuracy (Figure 2D labeled dot), but its TIGER rank is 5. This TF has five targets whose edges are all positive in both the prior network and “gold standard” set, meaning the prior edge sign is 100% accurate. During the course of Bayesian updating, TIGER wrongly flips two edges to negative. We further checked the edge weights and found that these two wrongly flipped edges have the lowest weights (Supplementary Figure S1), very close to zero, indicating that TIGER has low confidence in the sign of these edges. Therefore, when the prior network is very accurate, TIGER may make performance worse by trying to improve the network when the data and prior are not consistent with each other (Supplementary Figure S2). We implemented a simulation study to more precisely quantify this phenomenon. We started with a 100% accurate prior network, then sequentially flipped 10% more edge signs in each iteration and compared the performance degradation of VIPER and TIGER. When only ∼10-20% of edge signs are flipped, TIGER performs worse than VIPER. However, if >20% of edge signs are flipped, TIGER dominates.

### TIGER improves TF activity estimation by re-weighting edges

We next tested TIGER’s performance on two human cell lines. Both A375 melanoma and MCF7 breast cancer cells have TFKO datasets (T2), which have been used previously for TFA inference using the DoRothEA database [26]. DoRothEA is a collection of highly curated TF regulons that can serve as signed network priors for TF activity inference. We first compared TIGER and VIPER performance in TFA estimation (Figure 3A-D). Defining the top 5 ranked TFs as a “success” (due to the limited number of TFs in this data), TIGER again slightly outperformed VIPER on both datasets; however, the consistency between the methods was not as high as in the yeast dataset (Figure 3B, D). We reason this is because cancer cell lines have a larger genome and more complicated regulatory mechanisms. Additionally, the DoRothEA prior network is not specific to either of the A375 or MCF7 cell lines, meaning there could be many context-irrelevant edges. Thus, we hypothesized that TIGER’s better performance comes from its ability to re-weight edges, specifically, up-weighting true edges and down-weighting irrelevant edges. To test our hypothesis, we downloaded available ChIP-seq data for A375 and MCF7 from Cistrome DB [27]. For A375, there was ChIP-seq data available for 1 out of 14 knocked-out TFs, and for MCF7, there was data for 10 out of 25 knocked-out TFs. Among these 11 TFs, TIGER correctly identifies 7 of them in the top 5, but VIPER only identifies 4 of them (Figure 3E). TIGER seems to perform strictly better than VIPER because the 7 TFs TIGER identifies includes all 4 TFs that VIPER identifies.

**Figure 3.**
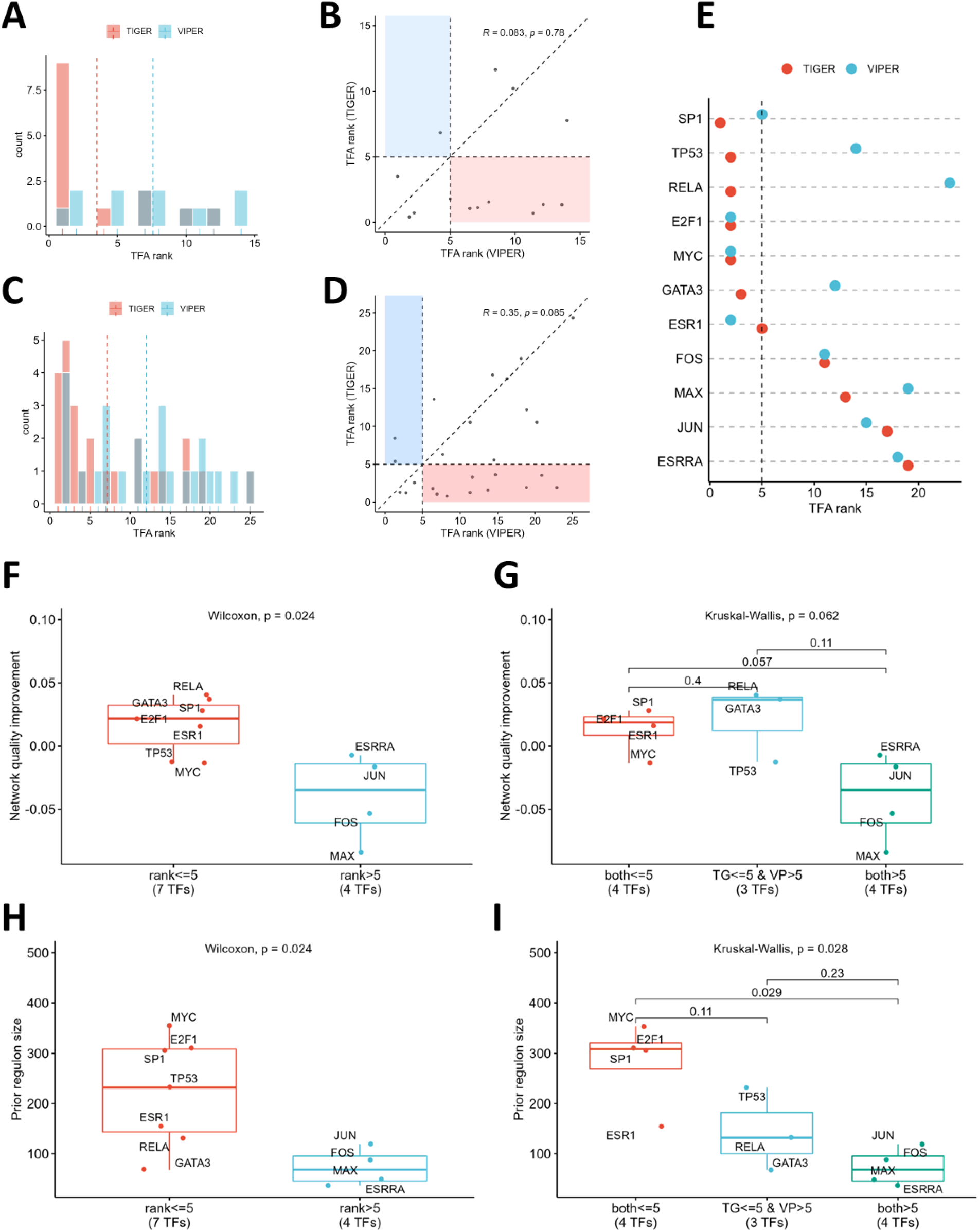
Comparison of TIGER and VIPER on cancer cell line TF knock-out datasets. (A-B). Overall comparison of TFA ranking in A375 TFKO data. A lower rank is better. (C-D). Overall comparison of TFA ranking in MCF7 TFKO data. A lower rank is better. (E). Dot charts of TIGER and VIPER’s performance on 11 TFs with ChIP-seq data. A lower rank is better. (F-G). Boxplot shows the association between the regulon Quality Score (QS) improvement (Y-axis) and TIGER’s performance (X-axis). TG=TIGER and VP=VIPER. (H-I). Boxplot shows the association between the regulon size (Y-axis) and TIGER’s performance (X-axis).

Using the ChIP-seq data, we asked whether TIGER’s performance is related to re-weighting edges. We first need to quantify the “correctness” of the edge weights. Thus, we defined a regulon quality score (QS), representing the inner product between the edge weights and ChIP-seq scores (see Methods). All edges have the same weight in the prior network, meaning that the regulon QS will be the arithmetic mean of all edges’ ChIP-seq scores. However, for the posterior network, all edges will be re-weighted; therefore, the posterior QS will be a weighted average of ChIP-seq scores. If TIGER correctly increases the weights of true edges (i.e., those with a higher ChIP-seq score) and decreases the weights of false edges (those with a lower ChIP-seq score), the posterior QS will be larger than the prior QS. With this definition, we found a significant correlation between TIGER’s success and regulon quality improvement (Figure 3F; Wilcoxon test, p<0.05).

Next, we asked if TIGER’s ability to improve QS leads to improved performance relative to VIPER. Out of the three TFs (RELA, GATA3, and TP53) on which TIGER has better performance, RELA and GATA3 have the largest QS improvement; however, TP53 has a decrease in the regulon quality (Figure 3G). As these three TFs have different behaviors, it appears that TIGER’s success is not driven by only one factor, i.e., regulon quality improvement. In addition to TP53, TIGER decreases the regulon quality of MYC, ESRRA, JUN, FOS, and MAX. Interestingly, both TIGER and VIPER could successfully identify MYC but failed to identify the other four TFs. How could TIGER fail to improve the network quality but still successfully identify MYC and TP53, and why is VIPER successful with only MYC? MYC and TP53 are well-studied cancer driver and suppressor genes; their DoRothEA prior regulon tends to be larger and more accurate due to the literature bias. This suggests that regulon size and quality could be influencing the performance of TIGER and VIPER.

To explore the effect of the prior regulon on algorithm performance, we first looked at regulon sizes across all TFs. Not surprisingly, MYC, TP53, and other TFs that were successfully identified by TIGER (i.e., among the top 5) have significantly larger regulon sizes than the four failed TFs (Figure 3H; Wilcoxon test, p<0.05). MYC has the largest regulon size (354 genes) and ESRRA has the smallest (37 genes). Both VIPER and TIGER are more successful at identifying TFs with large regulons and fail to identify very small regulons (Figure 3I; Wilcoxon test, p<0.05). However, TIGER has more tolerance for regulon size. For example, ESR1 has 155 putative targets, and TIGER and VIPER can both correctly identify it. RELA has 132 targets, but only TIGER can identify it. JUN has 119 targets, and both TIGER and VIPER failed to identify it. We also asked if performance could be related to prior regulon quality, but neither TIGER nor VIPER’s performance is correlated with the prior regulon quality (Supplementary Figure S3).

In conclusion, we found that TIGER can improve TFA estimation by improving edge weights in the regulatory network. TIGER and VIPER are both sensitive to prior regulon size, but TIGER is more tolerant to small regulons. We reason that this is because TIGER (1) uses a Bayesian framework to re-weight edges, thus filtering out noisy edges and (2) models the combined effect of multiple TFs on a target gene. When the regulon size is small, VIPER’s estimation can only use the information from a limited number of putative targets because each TF is considered completely independent, whereas TIGER can borrow information from other TFs.

### TIGER correctly identifies cell type-specific TF regulators

Since TIGER is able to refine the regulatory network to reflect context-dependent TFA, it is well-positioned to identify cell-type-specific regulatory mechanisms from scRNA-seq data (T3). We downloaded the PBMC10K scRNA-seq and scATAC-seq multiome dataset from the 10x Genomics website and used the Weighted Nearest Neighbor (WNN) method from “Seurat” [28] to integrate scRNA-seq and scATAC-seq data to cluster and identify cell types (Figure 4A). Then we used three methods to identify driver TFs for each cell type. The first is a multi-modality method, in which we combine motif scores computed from scATAC-seq using “Signac” [29] and “chromVAR” [29] with scRNA-seq information to estimate the most active TFs in each cell type. This method is regarded as the “gold standard” in this dataset because it incorporates scRNA-seq, scATAC-seq, and motif information. We then used this gold standard to evaluate the performance of TIGER and VIPER using only the scRNA-seq data and DoRothEA prior. We applied TIGER separately to each cell type, under the assumption that each cell type could have different regulatory mechanisms leading to cell-type-specific Bayesian updates on the GRN. On the other hand, we applied VIPER to each cell individually as it was designed to be used.

**Figure 4.**
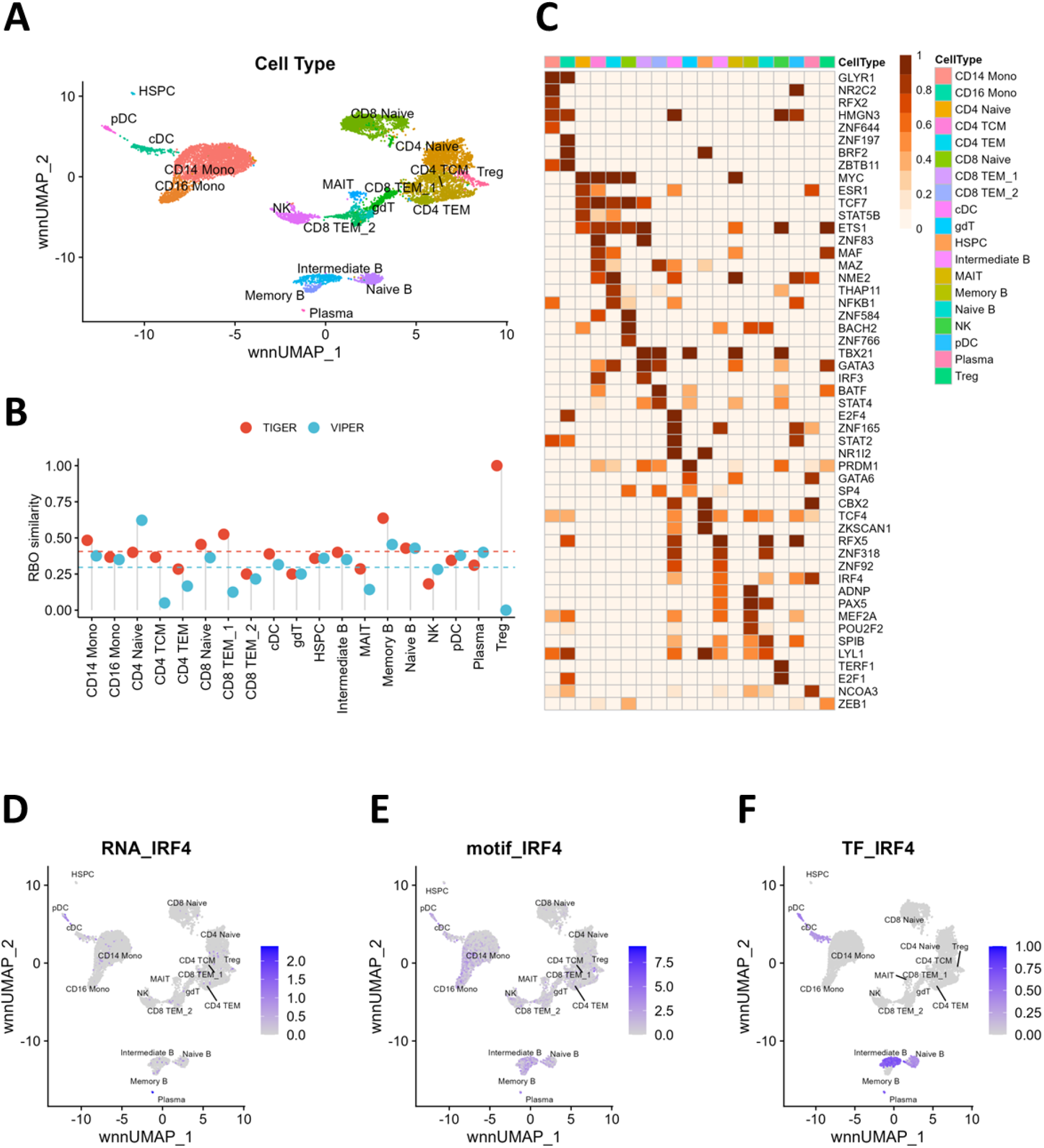
TIGER’s performance on PBMC scRNA-seq data. (A). UMAP plot of 19 cell types identified by the weighted nearest neighbor algorithm. (B). Comparison of TIGER and VIPER’s similarities with multi-modality analysis results in each cell type. Dashed lines are average similarity scores across all cell types. (C). Heatmap of activity scores of the top 5 TFs in each cell type. (D-F). Feature plots of IRF4 mRNA, motif, and TF activities. mRNA is the normalized gene expression score, the motif score is computed by combining scRNA-seq and scATAC-seq data, and the TFA score is estimated from TIGER.

To compare performance, we used the rank-biased-overlap (RBO) metric (see Methods) to quantify the similarity between multiple rankings of TFA. The RBO metric correlates two ranked vectors with more weight at the top of the ranked list where driver TFs should be found. Overall, the TIGER TF rankings had higher RBO than VIPER when compared with the multi-modality “gold standard” (Figure 4B; Wilcoxon Test, p<0.05). We visualized the top 5 TFs in each cell type using a heatmap (Figure 4C) and looked for evidence in the literature for their relevance to that cell type (Supplementary Table S1). For example, IRF4 is known to control differentiation of several types of Plasma, B cells, and DC cells but is not as critical for memory B cells [30-32]. Interestingly, mRNA levels of IRF4 indicate IRF4 has varying levels of expression in Plasma and some B and plasmacytoid dendritic cells (pDC) (Figure 4D). Multi-modality analysis shows IRF4 is enriched in Plasma, B cells, and DC cells, among others (Figure 4E). However, TIGER is the only method that reveals that IRF4 is the main driver in Plasma, Intermediate B, Naïve B, pDC, and conventional dendritic cells (cDC) but not in Memory B cells (Figure 4F) [32]. Moreover, TIGER is the only method that finds that PAX5 is the main regulator of Memory B cell differentiation [32] (Supplementary Figure S4). Similar phenomena are observed for TBX21 in NK cells [33], and for TCF4 in DC and monocytes [34, 35] (Supplementary Figure S5-S6). In general, the mRNA levels, multi-modality analysis, and TIGER results are consistent with each other, but TIGER may be able to uncover more subtle differences. Finally, we checked the top 20 target genes of IRF4 in intermediate B cells and compared the TIGER estimated edge weights to the IRF4 ChIP-seq score in B cells downloaded from Cistrome DB (Supplementary Figure S7). Though this analysis is not powered for significance, a positive Spearman correlation (R=0.25) suggests that TIGER may be able to model cell-type-specific regulatory mechanisms.

### TIGER outperforms other sign-constrained methods

VIPER can be used in conjunction with any TF regulons that the user selects. The previous results were all obtained with regulons from the Dorothea database, as these were demonstrated to provide the best performance. However, since TIGER is able to update the prior network and learn context-specific edge weights, we also compared TIGER’s performance with other methods. First, we used VIPER with data-driven TF regulons derived from ARACNe (VIPER-ARACNe) [3]. Second, we tried applying VIPER on the TIGER estimated network (VIPER-TIGER). Finally, since TIGER is based on matrix factorization (MF) method, we compared it with the only published MF method that incorporates sign constraints [18], which we call constrained MF (CMF). Using the same three TFKO datasets as above, we found that TIGER constantly outperforms all other methods (Figure 5A-C, Wilcoxon Test). Including context-specific regulons from ARACNe did improve the performance of VIPER on the two human cell line datasets but not as much in the yeast dataset, showing the importance of tissue-specific regulation in mammalian systems. We also observed that VIPER-TIGER has a better performance than VIPER and even works better than VIPER-ARACNe in two cases, which is consistent with the hypothesis that TIGER improves network quality. However, VIPER-TIGER does not have as stable performance across all TFs and datasets as TIGER. The constrained matrix factorization (CMF) method is slightly worse than TIGER, perhaps due to its inability to modify the prior edge signs.

**Figure 5.**
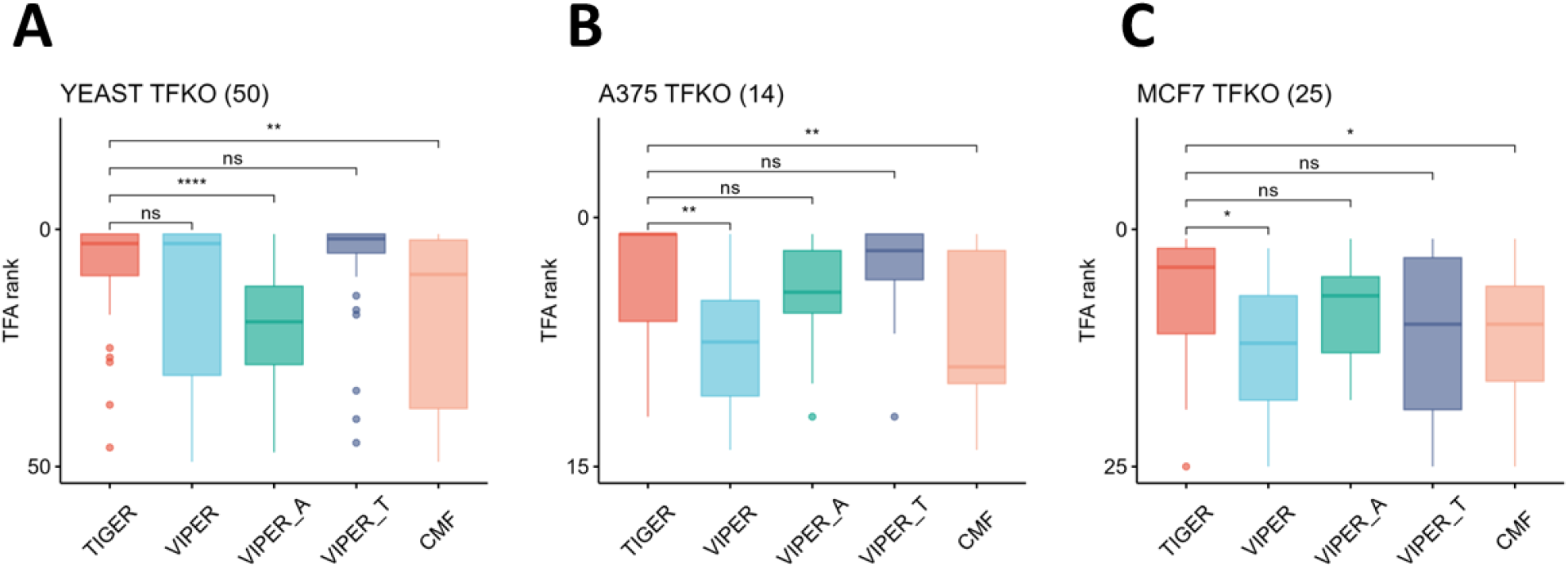
TIGER outperforms other sign-constrained TFA inference methods. (A-C) YEAST, A375, and MCF7 TFKO datasets. The Y-axis is the TFA ranking; lower rank (upper Y-axis) is better. The X-axis shows the five methods – TIGER, VIPER, VIPER_A, VIPER_T, and CMF. VIPER_A applies VIPER on the estimated ARACNe network. VIPER_T applies VIPER on the TIGER inferred network. CMF is the sign-constrained Matrix Factorization method. Pairwise comparisons between TIGER and other methods use Wilcoxon Rank Sum Test.

## Discussion

We present TIGER, a Bayesian matrix factorization method to better estimate regulatory networks and TF activity (TFA) in both bulk and single-cell RNA-seq data. Compared to other methods, TIGER has three main advantages. First, it distinguishes activation and inhibition events by incorporating prior edge signs and updating them with data. Second, it up-weights essential edges and shrinks irrelevant edges towards zero through a sparse Bayesian prior. Third, it uses a Variational Bayesian method to efficiently approximate the posterior distribution.

We found that TIGER outperforms multiple state-of-the-art TFA estimation methods in three validation datasets. VIPER is an efficient algorithm based on enrichment of TF regulomes among differentially expressed genes. VIPER can be applied to single samples and can correctly predict the driver TF when the prior network is accurate. In comparison, TIGER requires more samples as input but can use the data to refine the network and improve TFA estimation. TIGER performs better than VIPER on large yeast and mammalian datasets, regardless of whether VIPER is combined with static databases like DoRothEA or context-specific regulomes inferred using ARACNe. TIGER also performs better than a sign-constrained matrix factorization method. To explore what drives the improved performance of TIGER, we performed two types of analyses. In the yeast TFKO bulk RNA-seq data, we used ChIP data as a very accurate prior network structure and assessed TIGER’s ability to infer correct edge signs. In the cancer cell line TFKO bulk RNA-seq data, we used a less accurate, cell-type-agnostic prior network from DoRothEA to assess TIGER’s ability to refine edge weights. Our results indicate that edge sign and edge weight are both important for correct TFA estimation, and that edge sign has a higher impact – despite the fact that it is often ignored by network inference algorithms. TIGER is particularly useful when the prior edge sign accuracy is below 80%, because it can update and refine the signs accordingly. TIGER can also be applied in a cluster-by-cluster fashion to scRNA-seq data. Using the PBMC10k dataset, we found that this workflow improves identification of cell-type-specific regulators beyond what can be achieved using static regulome databases.

The performance of TIGER is limited by certain features of the data. In the yeast TFKO data, we noticed that TIGER’s Bayesian method attempts to update the network even if the prior network sign is 100% accurate, which can cause a decrease in TIGER’s performance. Although a fully accurate prior is not possible in reality, it would still be useful to minimize this issue. One could simply constrain the edge signs to be the same as the signs in the prior network, but this would damage performance when the prior edge signs are inaccurate. Another possibility is to construct a Bayesian credible interval for every edge. If the credible interval does not include zero, we have enough evidence to accept the edge sign (flipping); otherwise, the sign is unspecified, and perhaps the edge does not exist. An appropriate cutoff for the credible interval (e.g., 90% CI) could perhaps be determined by constraining the specific regulon sparsity and overall graph sparsity. We leave this for future study.

The TF regulon size can also affect the performance of TIGER. In the cancer cell line TFKO data, we observed that TFA estimation through TIGER and VIPER relies on having a reasonably large and accurate set of targets for each TF. This can be challenging for TFs that are not well-studied in the literature. One way to handle this problem may be to add computationally predicted edges into the prior network. However, this will increase computational time and decrease the quality of the prior network. Other possibilities are to use a different prior distribution or tune hyperparameters to control the network’s sparsity constraints.

The model framework underlying TIGER has several limitations. First, it does not account for non-linear TF-TF interactions arising from competitive or cooperative binding. Second, epigenetic effects on the susceptibility of each gene to regulation, for example, by making the promoter more or less accessible to TFs, are not considered. Thirdly, the linear model of TFA underlying TIGER cannot model saturation effects – i.e., when TFA increases past a certain threshold, the expression of the target gene may not change. In the future, we hope to incorporate TF-TF interactions by incorporating protein-protein interaction (PPI) information in the covariance of the TF-gene edge weight matrix *Z* (see Supplementary Information). To address the second problem, recent work [12] suggests that one could refine the prior networks by incorporating ATAC-seq data. For the third issue, we could use the logistic function to transform the linear combinations of TFA. The saturation of gene expression would then be reflected by the upper asymptotes. We leave these questions for future study.

In summary, we have developed a Bayesian matrix factorization method that takes as input multiple expression profiles and jointly infers context-specific regulomes and TF activity levels. When applied to bulk and single-cell RNA-seq data, TIGER outperforms other methods that cannot incorporate realistic biological features like sign of regulation or cell-type specificity. Future directions will continue to improve this method by integrating complementary data from burgeoning genomic technologies.

## Methods

### Testing Datasets

Yeast datasets were downloaded from [18], including TFKO, TFOE, and ChIP data. Expression data were already log-transformed and normalized, and we computed gene-wise Z-scores for input into TIGER and other methods. High-quality ChIP data were preselected based on the criterion in [18], yielding a network of 1104 edges between 50 TFs and 778 genes. Edge signs were inferred by comparing knock-out samples with wild-type samples.

A375 and MCF7 TFKO cell line data were downloaded from [26]. Data were already log-transformed and normalized, and we computed gene-wise Z-scores for input into TIGER and other methods. Low-quality TFKO samples were excluded from the analysis based on the criterion in [26], yielding 14 TFKO samples in A375 and 25 TFKO samples in MCF7. DoRothEA cancer prior network “dorothea_hs_pancancer” from “dorothea” R package 1.6.0 was used. We included all A-E level DoRothEA edges. These edges are curated from many resources, such as literature database, ChIP-seq, TFBS, TCGA inferred ARACNE [26]. ChIP-seq validation datasets for the 11 TFs were downloaded from Cistrome DB [27].

PBMC scRNA-seq and scATAC-seq data were downloaded from the 10X genomics website. Data preprocessing, weighted nearest neighbor integration, clustering, annotation, motif analysis, and driver TF identification follows the “Weighted Nearest Neighbor Analysis” vignette, updated on August 30, 2021. Specifically, R packages “Seurat” version 4.1.1, “Signac” version 1.7.0, “EnsDB.hsapiens.v86” version 2.99.0, “chromVAR” version 1.16.0, “JASPAR2020” version 0.99.10, “TFBSTools” version 1.32.0, “motifmatchr” version 1.16.0, “BSgenome.Hsapiens.UCSC.hg38” version 1.4.4, “presto” version 1.0.0 were used. TIGER and VIPER were applied on each cell type using “dorothea_hs” prior network from “dorothea” R package 1.6.0. Because scRNA-seq data is very sparse, in each cell type, we only considered the marker genes identified using the “presto” R package with a p-value cutoff of 0.01 and log-fold-change cutoff of 0, the same as in the vignette. IRF4 ChIP-seq data in B cells was also downloaded from Cistrome DB [27].

### TIGER

TIGER is based on Bayesian matrix factorization. Each gene expression sample *X*_*n*_ can be formulated as *X*_*n*_ = *WZ*_*n*_ + *∈*, where *W* is the regulatory network and *Z*_*n*_ is the vector of TF activities (TFA) in that sample. We use a Gaussian mixture prior to sparsely encode each element of *W* and a multivariate half-normal distribution for the non-negative TFA sample *Z*_*n*_. Parameter estimation is performed by Variational Bayes methods with a mean-field Gaussian distribution family, as implemented in the Automatic Differentiation Variational Inference (ADVI) algorithm in the probabilistic programming platform STAN [36]. The posterior mean is used to summarize results. Model checking is performed using cross-validation to estimate out-of-sample predictive accuracy. See Supplementary Information for a full description of the TIGER model and its implementation.

### VIPER

VIPER is a statistical test for the enrichment of each regulon based on the average ranks of the target genes [15]. It takes into account the sign of each TF-target interaction. We used the R package “VIPER” version 1.28.0. We used VIPER with default parameters, except the minimal regulon size filter is set to 0 to ensure no TFs will be removed in VIPER computation.

### Regulon Quality Score

Regulon Quality Score (QS) is defined as an inner product between edge weights and ChIP-seq scores.

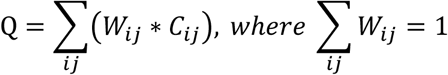

*W*_*ij*_ is the edge weight. For the prior network, *W*_*ij*_ = 1/*n. C*_*ij*_ is the ChIP-seq score, normalized to rank percentile. The prior Quality Score is the average ChIP-seq score of all targets. If TIGER up-weights the true edges and down-weights the irrelevant edges, posterior QS will increase. Thus, we defined the quality improvement as (Posterior QS – Prior QS)/Prior QS.

### RBO similarity

Rank-biased-overlap is used to evaluate two ranked lists based on the formula from “A Similarity Measure for Indefinite Rankings” [37]. Two ranking vectors with high RBO scores are similar especially at the top-ranked portions of the vectors. RBO ranges between 0 and 1. We used the R package “gespeR” version 1.26.0 to compute RBO similarity.

### ARACNE

ARACNE network was constructed using the ARACNE-AP algorithm [38]. One hundred steps of bootstrap were used with p-value<1E-8 as the cutoff.

### CMF

Constrained Matrix Factorization is a new variant of NCA that constrains the edge sign. We used the “TFAinference.py” python code v2.0.1 downloaded from [18]. Within 20 steps of iterations, the bilinear optimization algorithm converged in the three datasets we tested.

## Data and code access

All the testing datasets and analysis codes are available on Zenodo (https://doi.org/10.5281/zenodo.7425777). The TIGER package is available on GitHub (https://github.com/cchen22/TIGER).

## Acknowledgments

We acknowledge funding from NIH R01CA251729. We thank Jiawen Yang, Dante Bellomo, Akshita Sharma, and members of the Padi Lab for useful discussions.

## Author Contributions

C.C. conceptualized and developed the algorithms, designed the experiments, analyzed the data, and wrote the manuscript. M.P. directed the project, designed the experiments, and wrote the manuscript.

## Figure Legends

**Figure S1.**
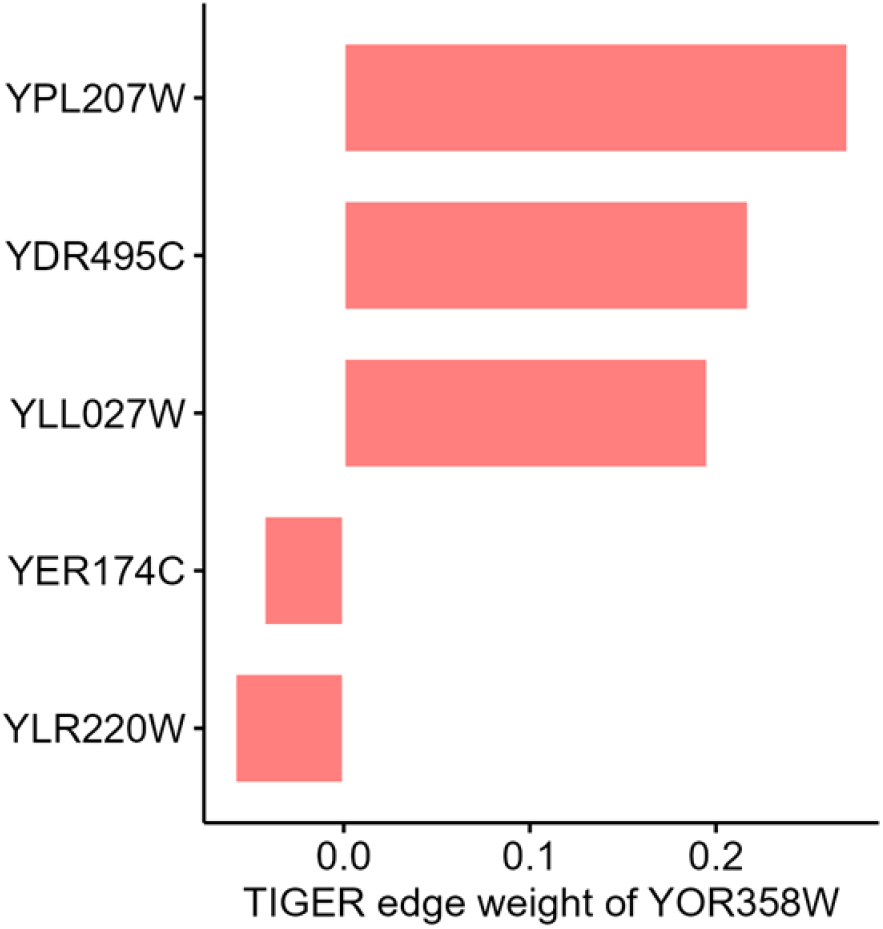
Five weighted targets of YOR358W.

**Figure S2.**
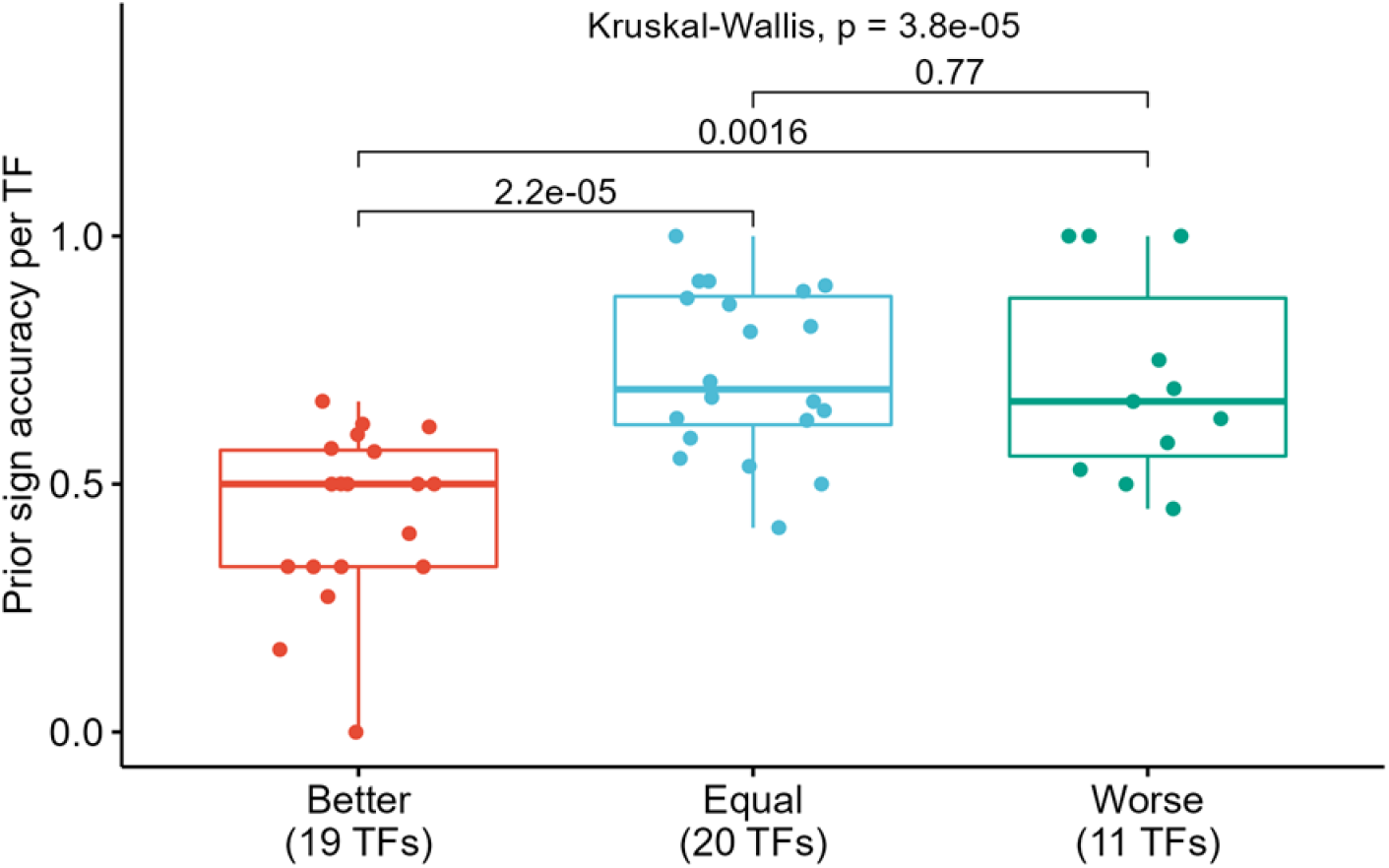
Prior sign accuracy in T1. For the 19 TFs where TIGER works better than VIPER, the prior sign accuracy is around 0.5, significantly lower than the other two clusters where VIPER has a good performance (KW test, p<0.05).

**Figure S3.**
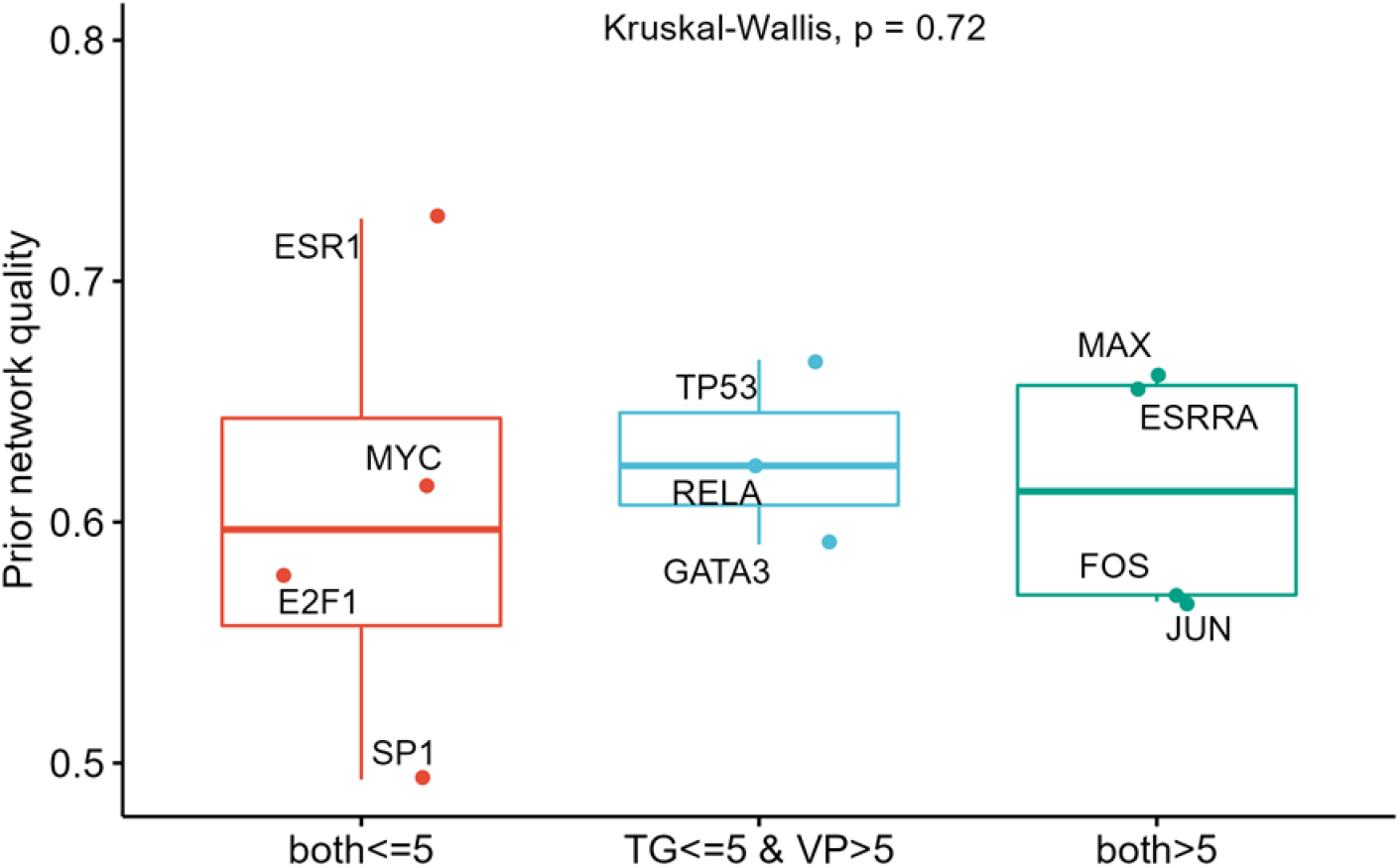
Prior regulon quality in T2. Regulon quality score (QS) does not have significant impact on TIGER or VIPER’s performance (KW test, p=0.72).

**Figure S4.**
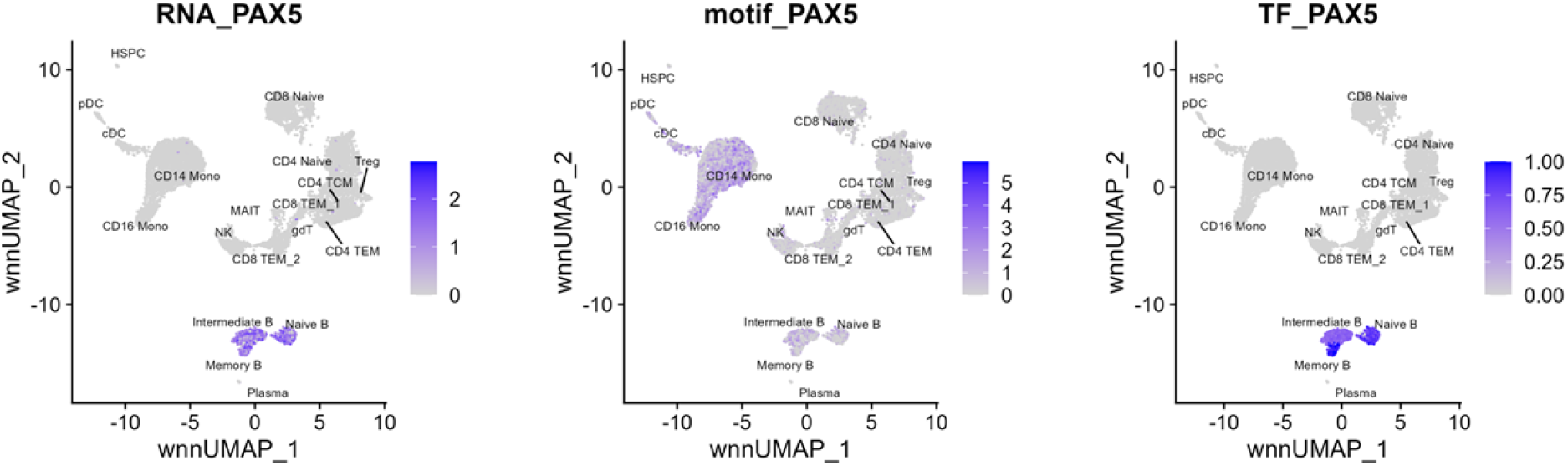
PAX5 mRNA expression, motif score, and TFA level in different cell types.

**Figure S5.**
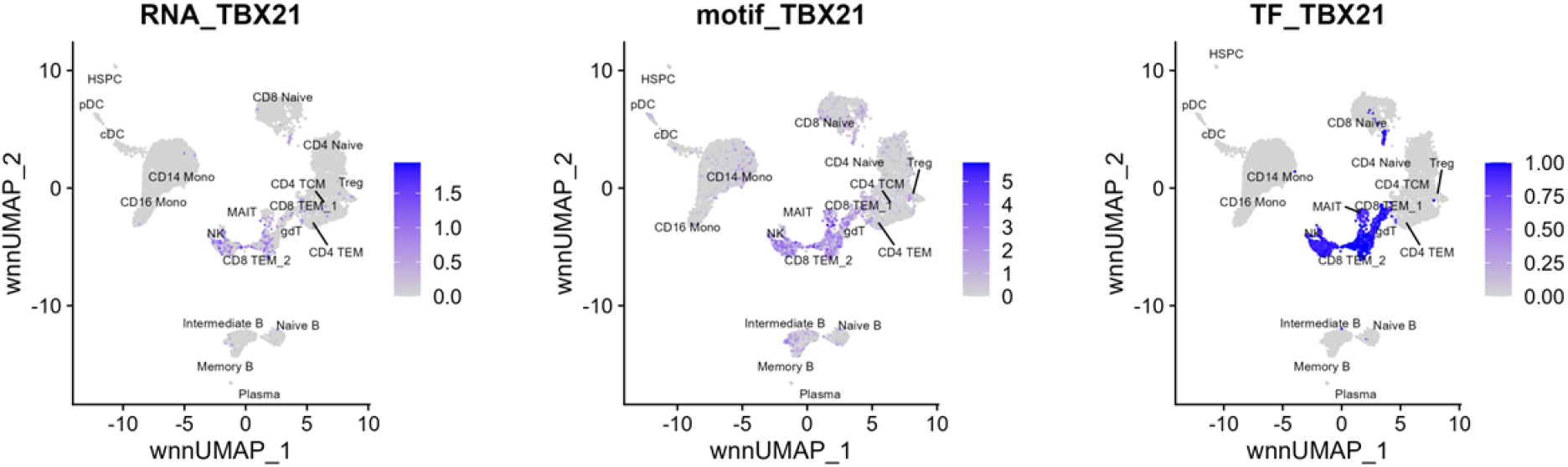
TBX21 mRNA expression, motif score, and TFA level in different cell types.

**Figure S6.**
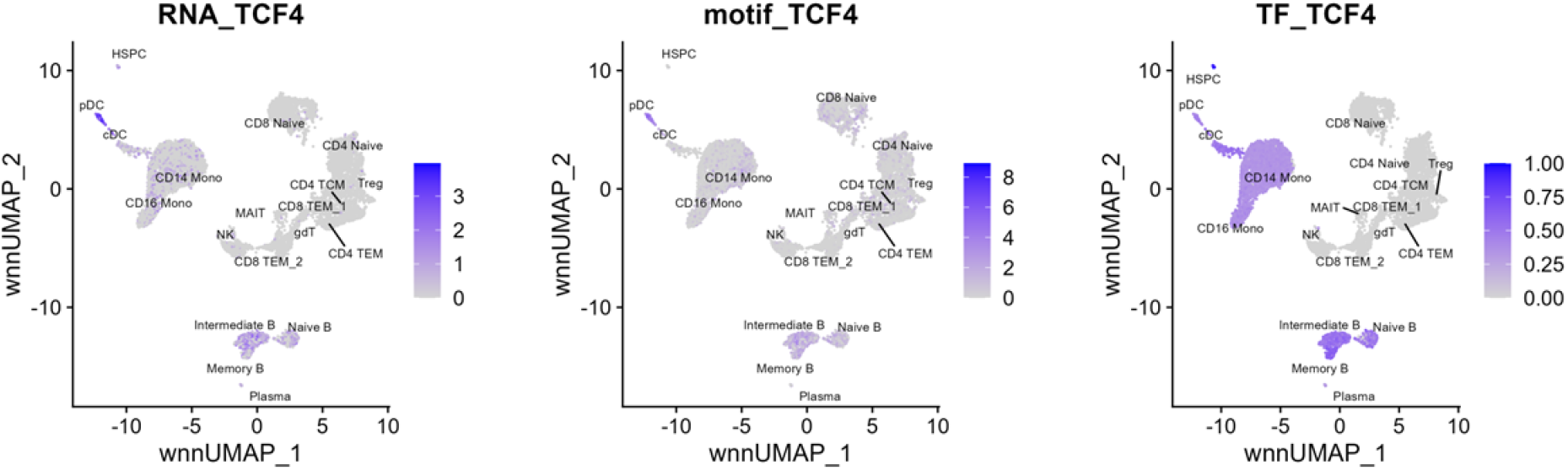
TCF4 mRNA expression, motif score, and TFA level in different cell types.

**Figure S7.**
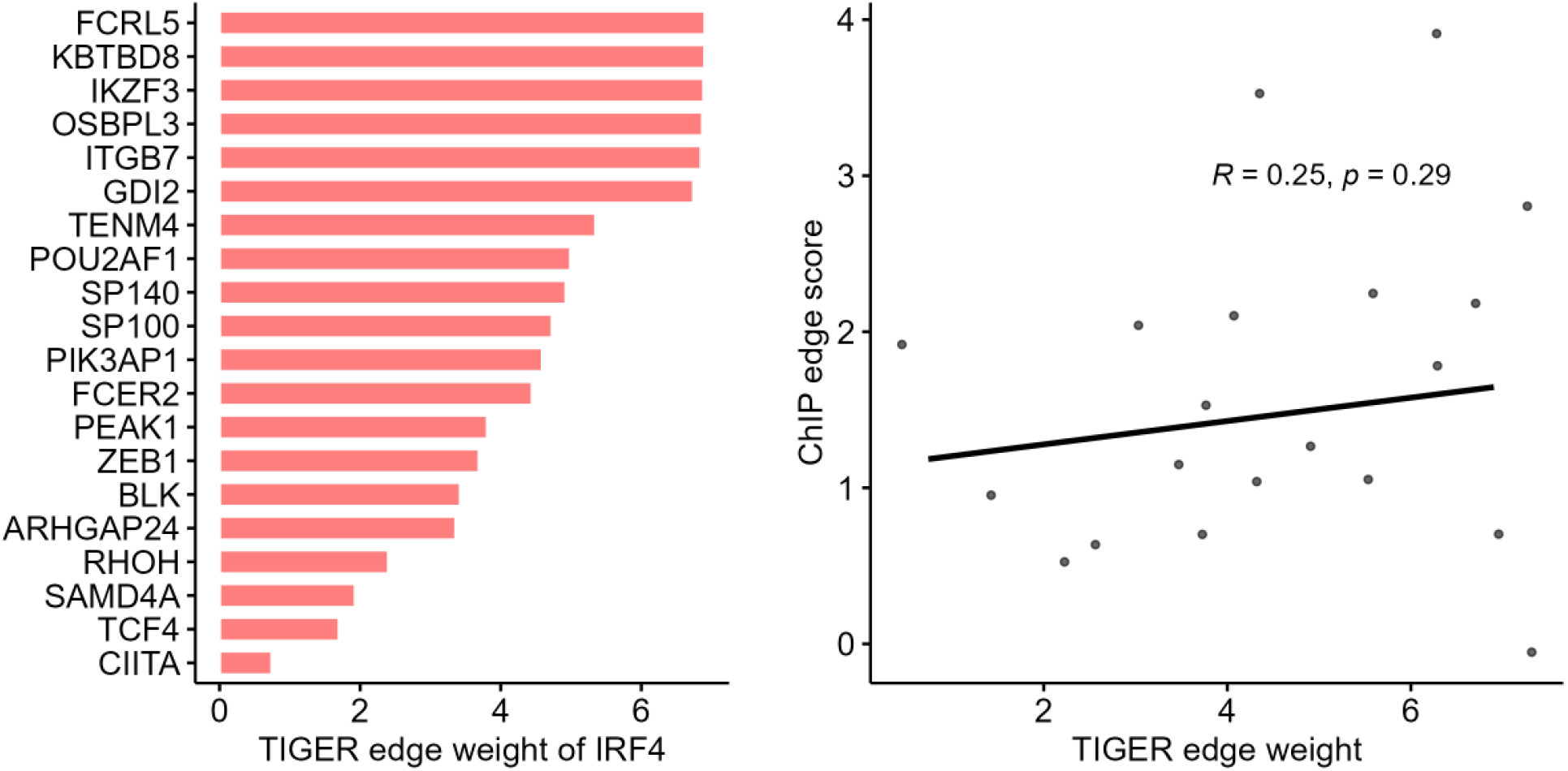
Top 20 weighted targets of IRF4 in intermediate B cells and their correlation with ChIP-seq target scores in B cells.

